# Macarthurs 1955 Stability Function is Related to Formal Dynamic Measures of Food Web Stability

**DOI:** 10.1101/477406

**Authors:** S. P. Keegan, S. E. Townsend, D.T. Haydon

## Abstract

Community complexity-stability relationships have been at the centre of ecological thinking for many decades. MacArthur (1955) proposed a measure of stability that reflected the diversity in the number of pathways energy can flow up through a food web but how this index correlates with more formal ideas of dynamical stability remains unexplored. Here, we examine the relationship between MacArthurs proposed index and measures of local and global stability in Lotka-Volterra food web models. Our results provide support for MacArthurs intuitive hypothesis that increasing the diversity of energy pathways through food webs endows them with greater stability, as measured by both the probability of local and global point stabilities, and the return time to stable equilibria following perturbation.

## Introduction

Sixty years ago Robert MacArthur set out a measure of community stability that he related to the fluctuations of animal populations (MacArthur, 1955). MacArthur viewed the food web as an energy transformer that could be modeled as an ergodic Markov chain, and he expressed a clear notion of the type of stability he had in mind: *”Suppose, for some reason, that one species has an abnormal abundance. Then we shall say the community is unstable if the other species change markedly in abundance as a result of the first. The less effect this abnormal abundance has on the other species, the more stable the community*.”

MacArthur’s 1955 paper has been heavily cited (1069 times at time of writing) and has contributed to a body of literature in support of the notion that ecological complexity would be positively related to community stability. This contrasts with the findings of a large number of papers demonstrating the opposite behavior in random model ecosystems (e.g. Gardner and Ashby 1970, May 1972, Daniels and MacKay 1974). We do not intend to review this lengthy and complicated debate, however it is an interesting historical anomaly that MacArthurs proposed measure of stability remains generally untested as a predictor of dynamical stability in model food webs (Rooney and McCann, 2012, but see Leigh, 1965).

MacArthur hypothesized that the amount of choice which energy has in following the paths up through a food web would be a measure of the stability of the community (*To see this, consider first a community in which one species is abnormally common. For this to have a small effect upon the rest of the community there should be a large number of predators among which to distribute the excess energy, and there should be a large number of prey species of the given species in order that none should he reduced too much in population*, MacArthur, 1955). His measure is more correctly viewed as one of trophic organization rather than stability per se (Horn, 1974).

## Methods

He proposed identifying all *K* energy pathways in a food web leading from basal species to top consumers, and calculating the proportion of the total energy, *p*_*w*_, that would reach the top consumer through the *w*th pathway. His measure of stability was the diversity of available pathways as captured by the Shannon-Weaver information index: 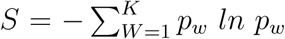. In a generalized Lotka-Volterra model defined in the standard way:

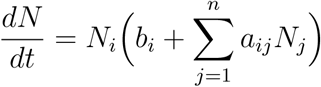

where *b*_*i*_ indicates the rate of change of species, density or biomass of the *i*th species in the absence of any other members of the community, and *a*_*ij*_ indicates the effect of species *j* on species *i*. The internal equilibrium point is given by *N*^∗^ = *−A*^−1^*b* (although for the equilibrium point to be ecologically plausible we require 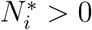 for all i). If j is a predator of i, the energy moving from i to j is proportional to 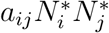. The proportion of energy, *q*_*ij*_, that *j* derives from *i* is 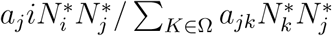 where Ω is the set of species from which *j* derives energy. *p*_*w*_ is then easily calculated as the product of all the *q*_*ij*_’s in an energy pathway from the bottom to the top of the food web (see Fig. 1 for a worked example). If more than one top consumer exists, we propose that the *p*_*w*_’s to each top consumer be weighted by the biomass of that top consumer divided by the sum of the biomass of all top consumers, thereby ensuring that 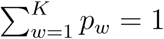 as originally proposed by MacArthur.

**Figure 1:**
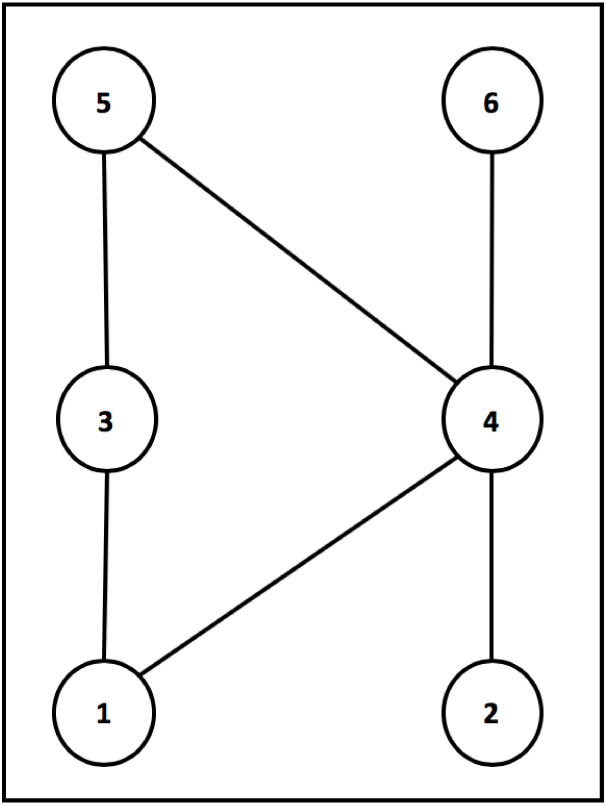
This food web has *K* = 5 energy pathways, p_1*−*3*−*5_, p_1*−*4*−*5_, p_2*−*4*−*5_, p_1*−*4*−*6_ and p_2*−*4*−*6_. Each of the *w* = 1..K unique pathways in the food web from autotroph to top predator is identified, and the product pw of the qijs contained in the path calculated (for example: 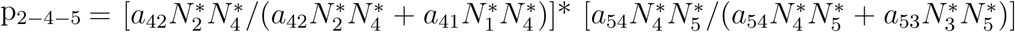 and weighted by the proportion of the biomass of top predators represented by the top predator that terminates each path. For example p_2*−*4*−*5_ would be weighted by 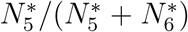 whereas p_2*−*4*−*6_ will be weighted by 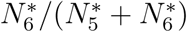.

Given a means of calculating MacArthurs index *S* for any Lotka-Volterra food web model, is it straightforward to relate *S* to stability measures of the model. Local stability can be assessed by inspection of the dominant eigenvalue of the Jacobian matrix **J** evaluated at the internal equilibrium point **N\*** (because at the internal equilibrium point 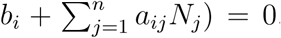, the *ij*th elements of which are given by 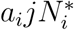).). This real part of the most positive eigenvalue, denoted *λ*_*d*_, must be negative if the equilibrium point is locally stable, and the return time of stable equilibria can be estimated as *−*1/*λ*_*d*_.

Local stability applies to only arbitrarily small perturbations and its ecological relevance is open to question. We can also determine whether the dynamics are permanent through an algorithm described by Law and Morton (1996). A dynamical system is permanent if no stable equilibrium points exist on its boundaries, and will therefore not experience extinction following a stochastic perturbation. Models with locally stable internal equilibrium points that are also permanent we term globally stable; models with locally stable internal equilibrium points that are not permanent we term fragile (a large enough perturbation may lead to extinction); models with locally unstable internal equilibrium points that are permanent we term oscillatory.

In this note, we describe how the stability properties of Lotka-Volterra model food webs relate to MacArthurs index *S*. Specifically we generate samples of food webs comprising different numbers of species with equal connectivity (defined as the proportion of *a*_*ij*_ elements (*i* ≠ *j*)not equal to zero), and examine how the proportion of locally stable (*P*_*LS*_), globally stable (*P*_*perm*_), fragile (*P*_*frag*_), and oscillatory food webs vary with *S*.

Food webs were constructed as follows: for a given level of connectivity (as close to 0.3 as possible) and number of species (*n*), trophic +/- links are randomly assigned in a manner that avoids the creation of loops of the form *a* preys on *b*, *b* preys on *c*, *c* preys on *a* (requiring the *a*_*ij*_ terms to be negative when *j > i*, and the corresponding *a*_*ij*_ term to be positive). Our algorithm then checks the resulting webs are at least minimally reticulated (no unconnected sub-webs), and calculates the number of autotrophs (defined as those species that have no predatory links to other species). Species that were not autotrophs were deemed to be heterotrophs. Interaction terms were assigned from uniform distributions as follows: autotroph self-regulatory terms U(0,-0.5); heterotroph self-regulatory terms U(0,1×10^*−*11^); the effects of predators on prey U(0,-0.5); and the effects of prey on predators U(0,0.2). The growth vector *b* was assigned values from a uniform distribution as follows: for autotrophs (0,1); for heterotrophs (0,-1). Food webs were deemed to be feasible if 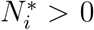 for all *i*. The Jacobian matrix (**J**) was then constructed from **A** and **N\***:

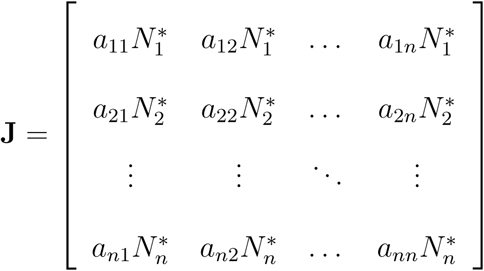

Food webs were generated that contained different numbers of species (ranging from n = 7 to 11). Generalized linear models were constructed to examine the relationship between measures of stability and MacArthur’s index. Simulations and analysis were performed in R (version 3.1.1).

Two thousand feasible food webs were generated for each community size (n=7 to 11). For each size, 50% of the sample was chosen to comprise locally stable food webs, and 50% locally unstable.Of these 10000 total food webs, 57% were permanent. Of these, 16% were oscillatory, and 9% fragile.

## Results

The probability of local stability (*P*_*LS*_) andMacArthurs index (*S*) were significantly positively related (*p <* 0.0001, *n* = 5000, GLM (binomial)), Fig. 2).Moreover, of food webs that were locally stable, we found a significant negative relationship between dominant eigenvalue (*λ*_*d*_) and MacArthurs index (*p <* 0.05 for all models, Fig. 3) indicating that the resilience of food webs with high values of MacArthurs index was greater (i.e. they had shorter return times).

**Figure 2:**
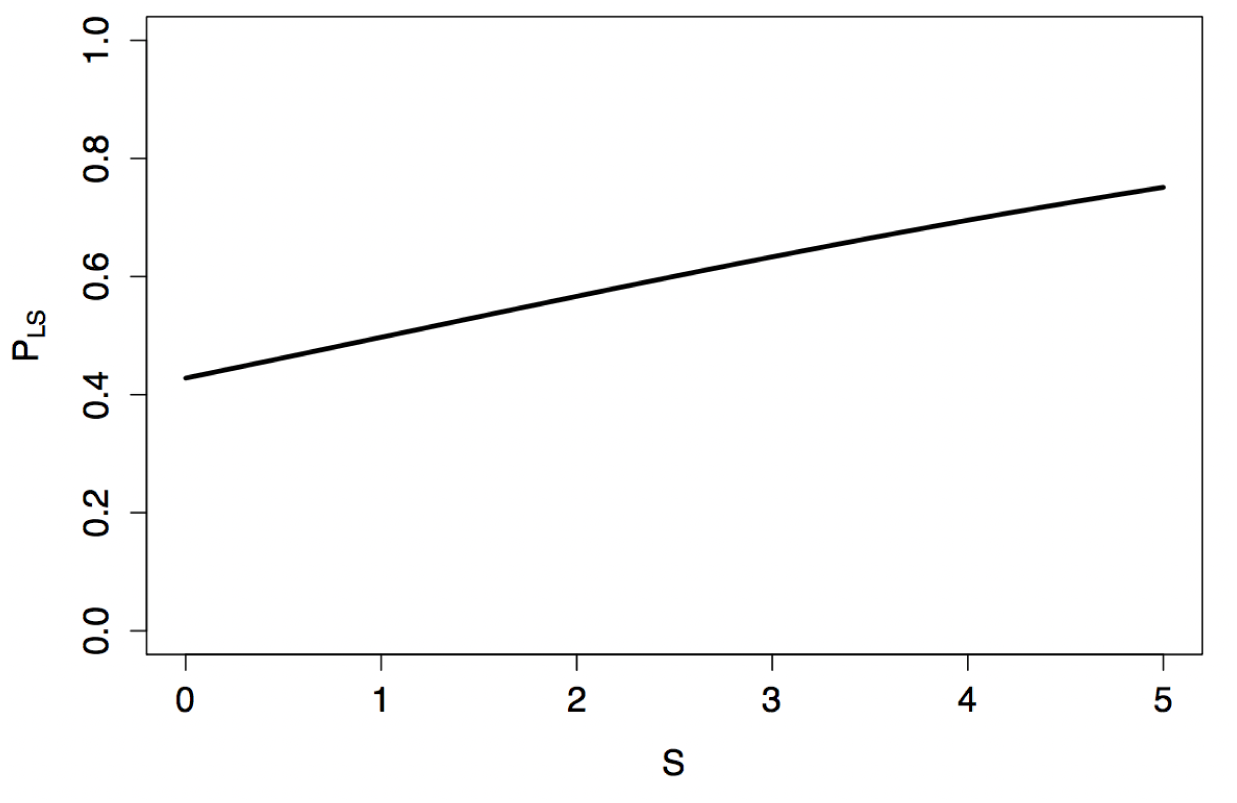
Probability of local stability predicted by a generalized linear model fitted to the results of simulations (*p <* 0.0001, *n* = 5000, GLM (binomial)). As MacArthurs index (*S*) increases, the probability that a food web will be stable also increases (all values of *n* combined).

**Figure 3:**
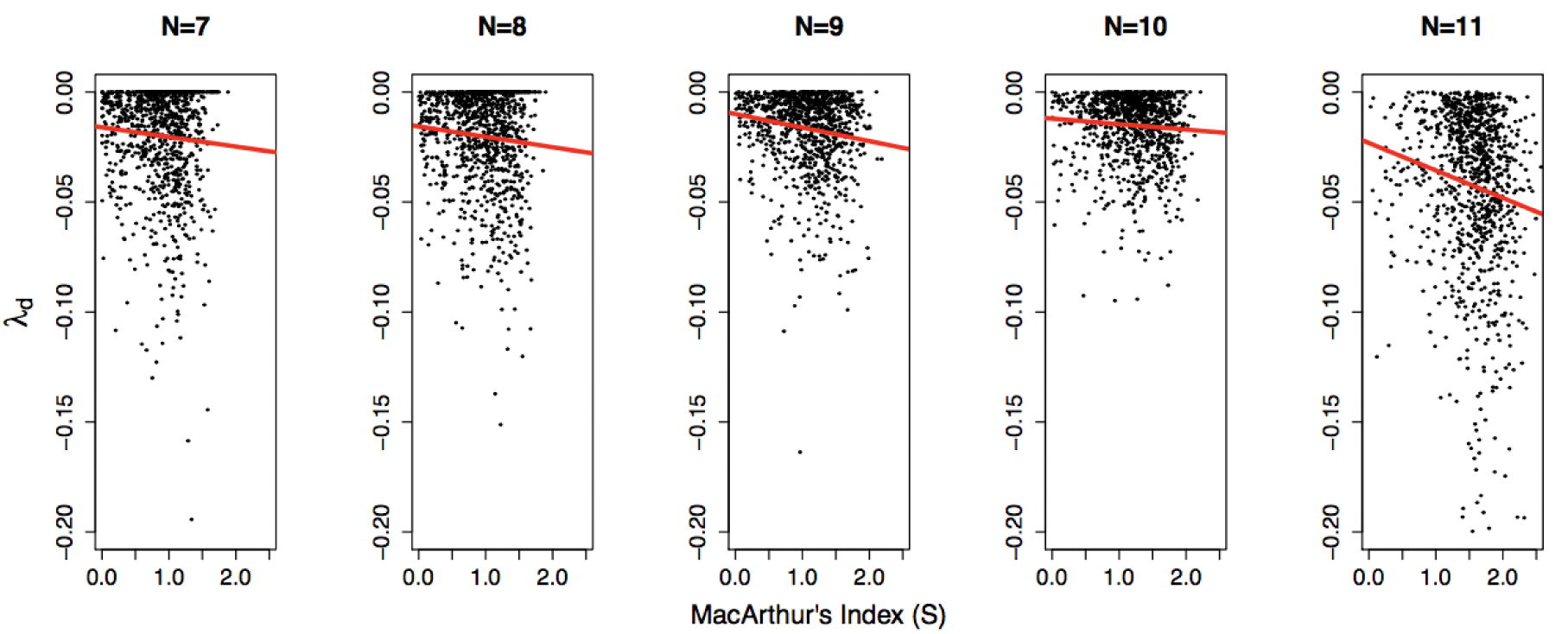
Dominant eigenvalue (*λ*_*d*_) with increasing MacArthurs index (*S*) and linear models plotted in red (*p*_*n*=7_ = 0.0234, *p*_*n*=8_ = 0.00598, *p*_*n*=9_ < 0.0001, *p*_*n*=10_ = 0.0218, *p*_*n*=11_ = 0.00154, *n* = 1000 for each sample, GLM). For each value of *n*, *λ*_*d*_ and *S* are negatively related. A more negative *λ*_*d*_ corresponds to a shorter return time to equilibrium.

We found no simple relationship between permanence andMacArthurs index, but we did find a significant interaction (*p <* 0.0001, *n* = 10000, GLM (binomial)) between number of species and MacArthurs indexsuch that food webs withfewer speciesshow a negative relationship with probability of permanence (*P*_*perm*_) as MacArthurs indexincreases (for *n ≤* 8), while for larger values of *n* (*n ≥* 9), the relationship is positive (Fig. 4).

**Figure 4:**
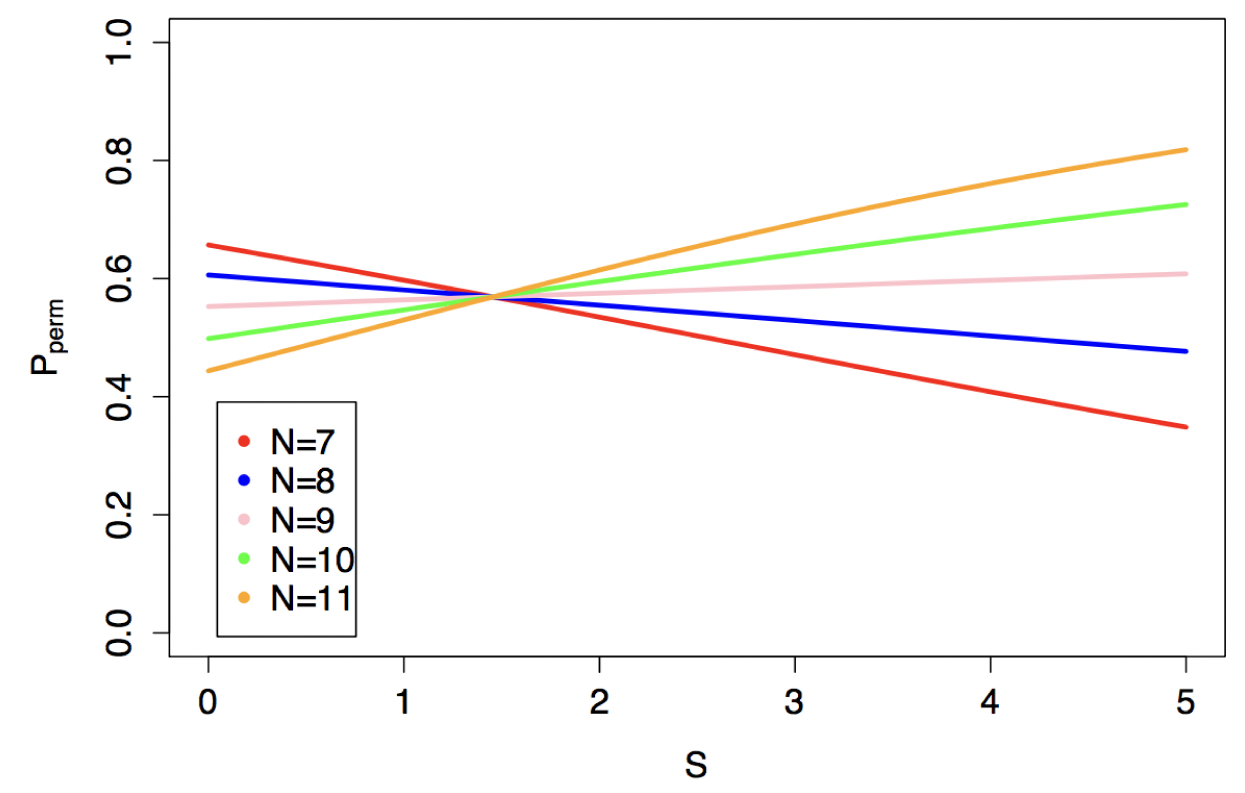
Probability of permanence as described by a generalized linear model. Interactions between MacArthurs index (*S*) and species number (*n*) (*p <* 0.0001, *n* = 10000, GLM (bino-mial)). For lower values (*n* = 7,8) as complexity in the form of *S* increases, the probability of permanence decreases. However, for higher values, as complexity increases, the probability of permanence increases.

For fragile webs (those that are locally stable but not permanent), there was also an interaction between MacArthurs index and number of species (*p <* 0.0001, *n* = 5000, GLM (binomial)). Smaller food webs (*n ≤* 10) showed an increase in the probability of fragility (Pfrag) asMacArthurs indexincreased whereas larger food webs (n = 11), showed a decrease in *P*_*frag*_ (Fig. 5). The probability of food webs being permanent but locally unstable did not vary significantly with MacArthurs index.

**Figure 5:**
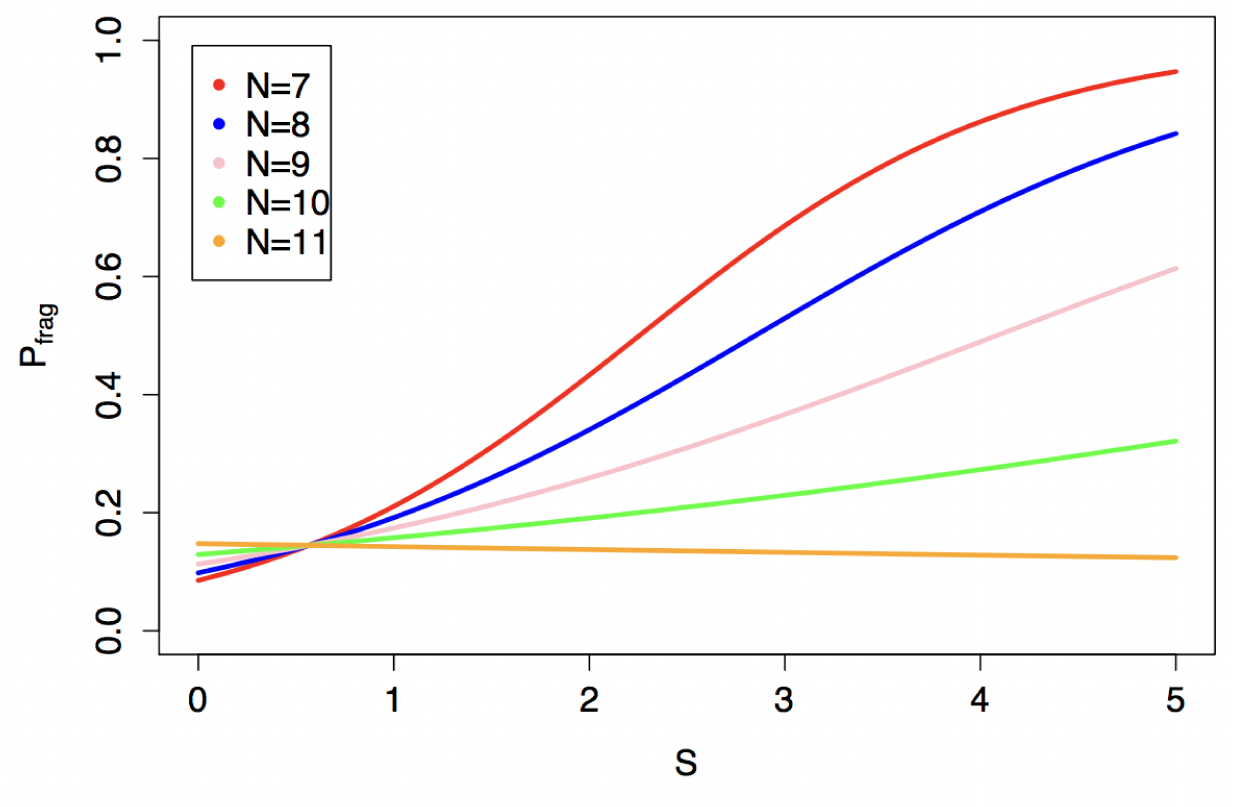
Probability of fragility as plotted by a generalized linear model. MacArthurs index (*S*) and species number (*n*) interact (*p <* 0.0001, *n* = 5000, GLM (binomial)). For higher values (*n* = 11) as complexity in the form of *S* increases, the probability of fragility decreases. However, for all lower values (*n* = 7-10), as complexity increases, the probability of fragility increases.

## Discussion

Our results indicate that MacArthurs proposed measure of stability predicts stability in much the same way as he anticipated: food webs with trophic organization corresponding to higher
 values of this measure are more likely to have equilibrium points that are locally stable, and when they are locally stable, they are likely to have faster return times to these equilibrium points following perturbation. Also in accordance with MacArthurs intuition, the frequency of oscillatory systems is not related to this measure.

MacArthurs 1955 study is prescient in a number of respects. First, in his anticipation of what type of stability future ecologists would devote so much effort to understanding, which is arguably close to measures of the time to return to some form of equilibrium following a perturbation. Second, in recognizing that *”stability can arise in two ways. First, it can be due to patterns of interaction between the species forming the community; second, it can be intrinsic to the individual species”*, thereby anticipating contemporary appreciation of the utility of separating the effects of the diagonal elements of the Jacobian matrix from its off-diagonal elements (e.g. Haydon 2000, Neutel et al 2002, Neutel et al 2007). And lastly, that the stability of food webs is likely to be governed by the subtle architecture of trophic structure, and not captured by simple measures of connectivity (Lawlor, 1978). For example, MacArthurs index appears to anticipate the importance of loop weight (Neutel et al 2002, Neutel et al 2007) and weak interactions (McCann et al 1998, Rooney and McCann, 2012) in determining the stability of food webs.

MacArthurs 1955 paper was an important early contribution to a very long-running area of ecological investigation. It was perhaps even more important than we thought.

## Acknowledgements

We thank Eric Pianka, Henry Horn and Mafalda Viana for helpful comments on the manuscript.

## References

Daniels, J. & MacKay, A.L., 1974. The stability of connected linear systems. Nature, 251, pp.4950

Gardner, M. R. & Ashby, W. R., 1970. Connectance of Large Dynamic (Cybernetic) Systens: Critical Values for Stability. Nature, 228, p.784

Horn, H. S., 1974. The Ecology of Succession. Annu. Rev. Ecol. Syst. 5:25–37

Law, R. & Morton, R. D., 1996. Permanence and the Assembly of Ecological Communities. Ecology, 77(3),pp.762–775

Lawlor, L. R., 1978. A Comment on Randomy Constructed Model Ecosystems. The American Naturalist, 112, pp.445–447

Leigh, E. G. 1965. On the Relation Between the Productivity, Biomass, Diversity, and Stability of a Community. Proc. Nat. Acad. Sci. 53, 777–783.

MacArthur, R., 1955. Fluctuations of Animal Populations and a Measure of Community Stability. Ecology, 36(3), pp.533536

May, R.M., 1972. Will a Large Complex System be Stable? Nature, 238, pp.413414

McCann, K., Hastings, A., & Huxel, G. R., 1998. Weak trophic interactions and the balance of nature. Nature, 395,pp. 794–798

Neutel, A., Heesterbeek, J. A. P., van de Koppel, J., Hoenderboom, G., Vos, A., Kaldeway, C., Berendse, F., & de Ruiter, P. C., 2007. Reconciling complexity with stability in naturally assembling food webs. Nature, 449, pp.559–602

Neutel, A., Heesterbeek, J. A. P., & de Ruiter, P. C., 2002. Stability in Real Food Webs: Weak Links in Long Loops. Science, 296,pp.1120–1123

Rooney, N., & McCann, K. S., 2012. Integrating food web diversity, structure and stability. Trends in Ecology and Evolution, 27(1),pp.40–46

